# High expression of JAM2 indicates better prognosis and immunotherapy response in breast cancer

**DOI:** 10.1101/2020.12.11.421081

**Authors:** Yang Peng, Chi Qu, Yingzi Zhang, Beige Zong, Bin Jian, Yong Fu, Jian Xie, Shengchun Liu

## Abstract

In our study, multiple databases were used to explore the potential role and underlying mechanism of junctional adhesion molecule B (JAM2) in breast cancer (BRCA). The data of JAM2 was downloaded from The Cancer Cell Line Encyclopedia (CCLE), the Genotype-Tissue Expression (GTEx), The Cancer Genome Atlas (TCGA) and Molecular Taxonomy of Breast Cancer International Consortium (METABRIC) databases. Receiver operating characteristic (ROC) curve analysis was performed to analyze the area under the curve (AUC) of JAM2 expression correlated with normal breast tissue and breast cancer tissue. Gene set enrichment analysis (GSEA) was used to identify the potential biological mechanisms of the JAM2. The expression of JAM2 mRNA were downregulated In most tumors, including BRCA, which may be due to the hypermethylated status. The AUCs, which were 0.929 and 0.887 by the logistic regression and random forest algorithms, indicated that JAM2 mRNA expression have good diagnostic value in BRCA. Univariate and multivariate analyses indicated JAM2 as an independent prognostic factor for the overall survival of BRCA patients in both the TCGA cohort (HR = 0.62, P = 0.034) and METABRIC cohort (HR = 0.77, P = 0.001). GSEA showed that multiple tumor pathways were suppressed in the JAM2 high expression group. The expression of JAM2 was most positively related to the epithelial-mesenchymal transition (EMT) score (r = 0.38; P <0.01) by the reverse-phase protein array (RPPA) analysis. Patients with high JAM2 expression may be more sensitive to immunotherapy. 18 chemotherapy drugs that patients in the JAM2 low expression group were more sensitive to were identified. Our results demonstrated the diagnostic and prognostic value of JAM2. Analysis of the molecular mechanisms indicates the potential role of JAM2 as a tumor suppressor, and high JAM2 expression may predict a better immunotherapy response in BRCA.

## Introduction

Breast cancer (BRCA), one of the most prevalent malignancies in women, is one of the most common cancers worldwide along with lung and colon carcinoma and primarily contributes to malignancy-related mortality in women[1, 2]. In 2020, approximately 276,480 new cases of BRCA will be diagnosed among women in the US, and 42,170 of these women will consequently die from it[3]. Mortality from BRCA has continuously decreased since 1989 due to improvements in treatment and early detection by mammography[4]. However, not all patients have benefitted uniformly from these advances, as mostly indicated by the distinct divergence in the application of adjuvant hormonal therapy due to the versatile classification of BRCA. Currently, BRCA is dominantly divided into 4 distinct subtypes according to the status of estrogen receptor (ER), progesterone receptor (PR), human epidermal growth factor receptor 2 (HER2) and Ki-67, and the treatment regimens for different subtypes vary dramatically[5]. Although advances have been dedicated to therapeutic regimens and the discovery of new drugs, the outcomes of BRCA patients with metastasis remain obstinately poor and have hardly improved. In fact, distant metastasis is responsible for over 90% of BRCA-related mortality[6]; hence, identifying metastatic targets for BRCA is of great interest and is essential.

Junctional adhesion molecule B (JAM2), a member of the immunoglobulin superfamily (IgSF), is located on the membrane of eukaryotic cells and enriched at cell junctions in epithelial and endothelial cells[7]. Based on its biofunction, JAM2 has been reported to participate in multiple biological processes, including tight junction assembly, regulation of paracellular permeability, angiogenesis, tumor metastasis and cell proliferation[8–10]. Structurally distinguished from other JAM family members, JAM2 could specifically interact with JAM-C and integrin α4β1 and subsequently form multimer interactions with integrin counter-receptors[11]. Indeed, it was reported that diminishing the interactions of JAM-B and JAM2 significantly reduced monocyte numbers in the abluminal compartment by increasing reverse transmigration[12]. Moreover, Ebnet et al identified the interaction between PAR-3 and the JAM2 and JAM-C complex at its first PDZ domain[13]. Additionally, JAM2 was reported to negatively regulate pro-angiogenic pathways by inhibiting VEGF-induced ERK1/2 activation[14]. Importantly, JAM2 has been demonstrated to play an important role in the metastasis of multiple tumor cells. Hajjari et al showed that the JAM2 and JAM-C complex could activate the expression of the actin filament-associated protein (AFAP) gene and that the dysregulation of JAM2 and JAM-C potentially contributed to the progression of gastric adenocarcinoma tumors[15]. In addition, the interaction of JAM2 and JAM-C could activate c-Src, a proto-oncogene, and therefore regulate cell invasion and migration in glioma[16]. Interestingly, overexpression of JAM2 inhibited VEGF-induced angiogenesis and thus attenuated the progression of lung carcinoma in vivo[17]. However, the expression levels of JAM2 vary dramatically among different tumor types, and the role of JAM2 in breast cancer has rarely been evaluated.

## Materials and Methods

### Data collection

We downloaded 21 tumor cell lines from the Cancer Cell Line Encyclopedia (CCLE) database (https://portals.broadinstitute.org/ccle/data) and 31 normal tissues from the Genotype-Tissue Expression (GTEx) database (http://www.gtexportal.org). The pan-cancer‐normalized data of 27 tumor and normal tissues were downloaded from UCSC Xena (https://tcga.xenahubs.net). The clinical and RNA sequencing (RNA-seq) data of 947 patients in The Cancer Genome Atlas (TCGA) cohort and 1891 patients in the Molecular Taxonomy of Breast Cancer International Consortium (METABRIC) cohort of BRCA were obtained from TCGA portal (https://tcga-data.nci.nih.gov/tcga/) and cBioPortal (http://www.cbioportal.org). Data on the methylation status of BRCA were extracted from the FireBrowse database (http://www.firebrowse.org). A total of 694 oncogenes were obtained from the ONGene database[18] (http://ongene.bioinfo-minzhao.org/), and 589 tumor suppressor genes (TSGs) were downloaded from the TSGene 2.0 database[19] (https://bioinfo.uth.edu/TSGene/). All data are publicly available.

### Gene set enrichment analysis (GSEA)

The BRCA samples were separated into high and low JAM2 expression groups based on the optimal cutoff value of JAM2 by the surv_cutpoint function of the survminer R package (https://CRAN.R-project.org/package=survminer). The clusterProfiler R package was used to perform GSEA[20]. The annotated gene set files “c2.cp.kegg.v7.1.entrez.gmt” and “h.all.v7.1.entrez.gmt” were used as references. The significance threshold was set as p value < 0.05.

### Estimation of immunotherapy and chemotherapy response

The estimated scores, immune scores and stromal scores of BRCA were analyzed by the *estimate* R package[21]. The Immune Cell Abundance Identifier (ImmuCellAI) (http://bioinfo.life.hust.edu.cn/web/ImmuCellAI/), which uses a gene set signature-based method to precisely estimate the infiltration score of 24 immune cell types, was used to predict the immunotherapy response (anti-PD1 or anti-CTLA4 therapy) of BRCA in the TCGA and METABRIC cohorts[22]. The chemotherapy response was estimated by the half maximal inhibitory concentration (IC50) of each BRCA patient from the TCGA cohort through the *pRRophetic* R package[23].

### Statistical analysis

All statistical analyses were performed using R version 4.0.0 (2020-04-24). For continuous variables, the Kruskal-Wallis H test was performed. Univariate and multivariate Cox regression analyses were used to identify the predominant prognostic factors of overall survival (OS) (P < 0.05). Kaplan-Meier survival curves were compared using the log-rank test. The ggplot2 R package (https://CRAN.R-project.org/package=ggplot2) was used to plot the correlation bubble diagram. P < 0.05 (two-sided) was considered statistically significant.

## Results

### The mRNA expression of JAM2 from pan-cancer analyses of different databases

Three different databases were used to explore the mRNA expression of JAM2 in different tumors and normal tissues. First, the data of each tumor cell line were downloaded from the CCLE database, and the expression of JAM2 was observed in 21 different tissues (Fig 1A). In addition, the GTEx database was used to analyze the expression levels of JAM2 in 31 normal tissues, as shown in Fig 1B. Interestingly, compared to that in other tumor cells, the expression of JAM2 in BRCA cell lines was relatively low, but compared to that in other normal tissues, the expression of JAM2 in BRCA cells was relatively high. Considering the small number of normal samples in TCGA, we integrated the normal tissues from the GTEx database into the TCGA database to analyze the differential expression of JAM2 in 27 tumor and normal tissues (Fig 1C). Except for six tumors (marked in blue), the expression of JAM2 was low in all other tumors (marked in yellow) compared to normal tissues.

**Fig 1.**
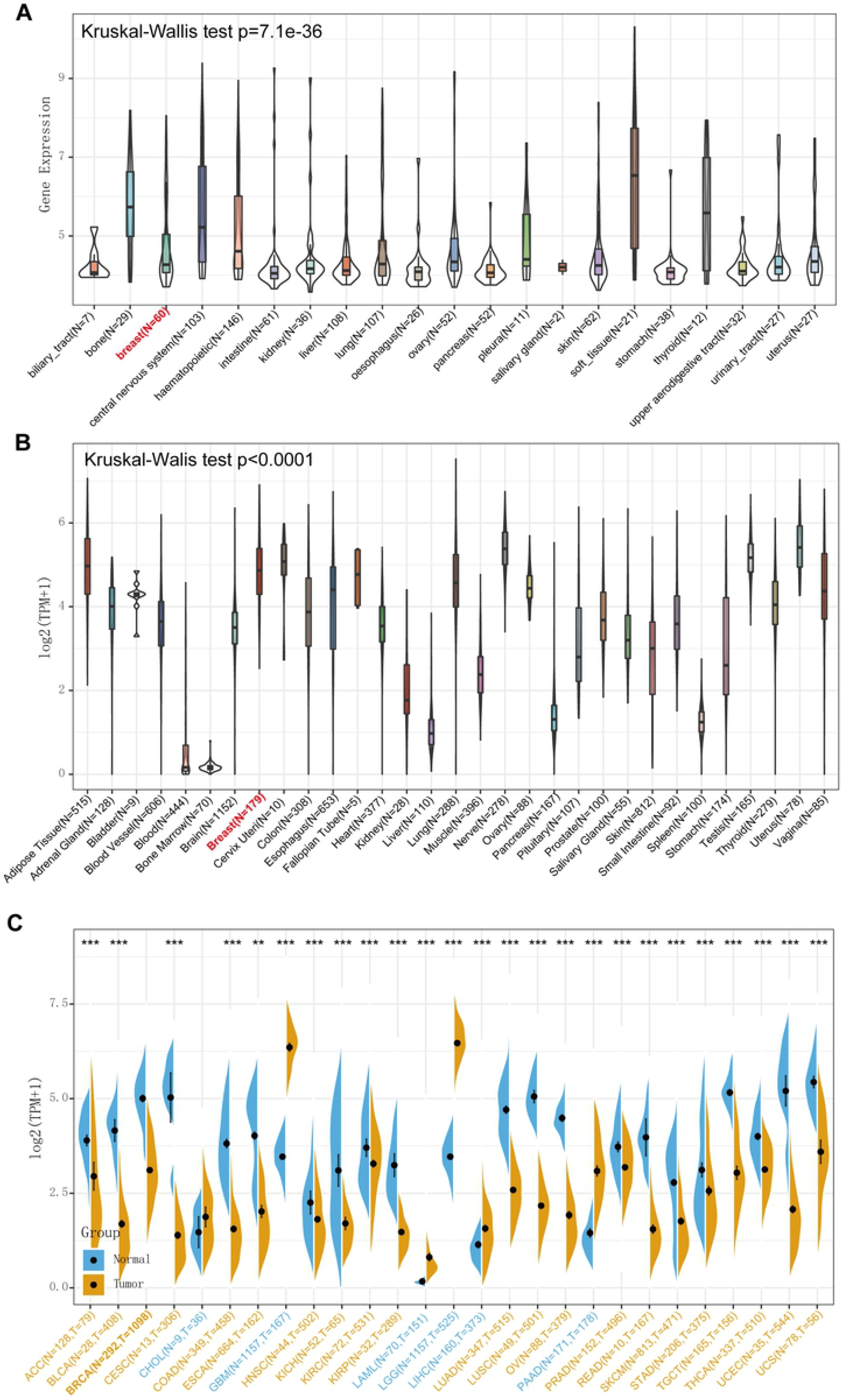
The mRNA expression of JAM2 from pan-cancer analyses of different databases. The mRNA expression of JAM2 in 33 normal (GTEx pan-normal) tissues (A); the red marker is normal breast tissue (n = 60). The mRNA expression of JAM2 in 21 tumor (CCLE pan-cancer) cells (B); the red markers are breast cancer cells (n = 179). The differential expression of JAM2 in 27 tumor samples (TCGA tumor vs TCGA normal + GTEx normal); yellow markers indicate low expression of JAM2 in these tumors, and blue markers indicate the opposite. The Kruskal–Wallis test was performed to calculate the P-value, and those associations with a P-value < 0.01 were considered significant.

### The diagnostic and prognostic value of JAM2

To evaluate the mRNA expression level of JAM2 in human BRCA, two independent datasets were used for analysis. The clinical data of the 947 patients in the TCGA cohort and 1891 patients in the METABRIC cohort of BRCA are supplied in table 1 and table 2 (Table S1). The results indicated that the expression of JAM2 was downregulated in BRCA tissues compared to normal tissues in the TCGA cohort (Fig 2A). Moreover, the expression of JAM2 was altered in subtypes of breast cancer; among them, it was highly expressed in the claudin-low subtype and expressed at low levels in the basal subtype (Fig 2B). The same result was confirmed in the METABRIC database (Fig 2C, 2D). In addition, to identify the diagnostic and prognostic value of JAM2 in BRCA, receiver operating characteristic (ROC) curve analysis was performed to analyze the area under the curve (AUC) of JAM2 expression correlated with normal breast tissue and breast cancer tissue by two algorithms. As shown in Fig 2E, the AUCs were 0.929 and 0.887 by the logistic regression and random forest algorithms, respectively, in the TCGA cohort, indicating that JAM2 has good clinical diagnostic value. Furthermore, the patients were separated into either the JAM2 high expression group or the JAM2 low expression group based on the optimal cutoff value, which was derived from the *surv_cutpoint* function of the *survminer* R package. The results indicated that patients with low JAM2 mRNA expression had a lower OS rate (Fig 2F; P = 0.016). Next, the METABRIC dataset was used to validate the results, and the same results were produced (Fig 2G, 2H).

**Table 1.** Correlation between the clinicopathologic variables and JAM2 mRNA expression in breast cancer in TCGA cohort.

| Parameters | High (N=256) | Low (N=691) | P-value |
| --- | --- | --- | --- |
| **Age** |  |  |  |
| Mean (SD) | 56.6 (12.4) | 59.1 (13.4) | 0.00806 |
| Median [Min, Max] | 56.0 [26.0, 90.0] | 59.0 [26.0, 90.0] |  |
| **Menopausal State** |  |  |  |
| Post | 151 (59.0%) | 483 (69.9%) | 0.00198 |
| Pre | 105 (41.0%) | 208 (30.1%) |  |
| **Histologic Subtype** |  |  |  |
| Ductal/NST | 134 (52.3%) | 549 (79.5%) | <0.001 |
| Other | 122 (47.7%) | 142 (20.5%) |  |
| **Histologic Grade** |  |  |  |
| NA | 256 (100%) | 691 (100%) | <0.001 |
| **Tumor Stage** |  |  |  |
| I | 53 (20.7%) | 109 (15.8%) | 0.109 |
| II | 137 (53.5%) | 415 (60.1%) |  |
| III | 64 (25.0%) | 154 (22.3%) |  |
| IV | 2 (0.8%) | 13 (1.9%) |  |
| **ER Status** |  |  |  |
| Negative | 21 (8.2%) | 171 (24.7%) | <0.001 |
| Positive | 227 (88.7%) | 491 (71.1%) |  |
| Missing | 8 (3.1%) | 29 (4.2%) |  |
| **PR Status** |  |  |  |
| Negative | 45 (17.6%) | 237 (34.3%) | <0.001 |
| Positive | 203 (79.3%) | 424 (61.4%) |  |
| Missing | 8 (3.1%) | 30 (4.3%) |  |
| **HER2 Status** |  |  |  |
| Negative | 189 (73.8%) | 485 (70.2%) | 0.104 |
| Positive | 30 (11.7%) | 113 (16.4%) |  |
| Missing | 37 (14.5%) | 93 (13.5%) |  |
| **Overall survival (month)** |  |  |  |
| Mean (SD) | 42.1 (29.4) | 35.3 (27.1) | 0.00134 |
| Median [Min, Max] | 33.6 [2.13, 120] | 25.6 [1.03, 117] |  |
| **Vital status** |  |  |  |
| Living | 228 (89.1%) | 596 (86.3%) | 0.301 |
| Deceased | 28 (10.9%) | 95 (13.7%) |  |

**Notes:** \*P < 0.05.
**Abbreviations:** JAM2, The junctional adhesion molecule B; TCGA, The Cancer Genome Atlas;
ER, estrogen receptor; PR, progesterone receptor; HER2, human epidermal growth factor receptor 2.

**Table 2.** Correlation between the clinicopathologic variables and JAM2 mRNA expression in breast cancer in METABRIC cohort.

| Parameters | High (N=595) | Low (N=1296) | P-value |
| --- | --- | --- | --- |
| **Age** |  |  |  |
| Mean (SD) | 58.4 (12.7) | 62.3 (12.9) | <0.001* |
| Median [Min, Max] | 58.7 [21.9, 89.1] | 63.3 [26.7, 96.3] |  |
| **Menopausal State** |  |  |  |
| Post | 430 (72.3%) | 1051 (81.1%) | <0.001* |
| Pre | 165 (27.7%) | 245 (18.9%) |  |
| **Histologic Subtype** |  |  |  |
| Ductal/NST | 398 (66.9%) | 1046 (80.7%) | <0.001* |
| Other | 195 (32.8%) | 238 (18.4%) |  |
| Missing | 2 (0.3%) | 12 (0.9%) |  |
| **Histologic Grade** |  |  |  |
| 1 | 82 (13.8%) | 82 (6.3%) | <0.001* |
| 2 | 260 (43.7%) | 475 (36.7%) |  |
| 3 | 228 (38.3%) | 697 (53.8%) |  |
| Missing | 25 (4.2%) | 42 (3.2%) |  |
| **Tumor Stage** |  |  |  |
| I | 193 (32.4%) | 281 (21.7%) | 0.00106* |
| II | 237 (39.8%) | 560 (43.2%) |  |
| III | 41 (6.9%) | 74 (5.7%) |  |
| IV | 3 (0.5%) | 6 (0.5%) |  |
| Missing | 121 (20.3%) | 375 (28.9%) |  |
| **ER Status** |  |  |  |
| Negative | 113 (19.0%) | 328 (25.3%) | 0.0031* |
| Positive | 482 (81.0%) | 968 (74.7%) |  |
| **PR Status** |  |  |  |
| Negative | 256 (43.0%) | 633 (48.8%) | 0.0212* |
| Positive | 339 (57.0%) | 663 (51.2%) |  |
| **HER2 Status** |  |  |  |
| Negative | 536 (90.1%) | 1121 (86.5%) | 0.0336* |
| Positive | 59 (9.9%) | 175 (13.5%) |  |
| **Overall survival (month)** |  |  |  |
| Mean (SD) | 132 (74.6) | 122 (76.9) | 0.0142* |
| Median [Min, Max] | 122 [2.30, 301] | 114 [1.23, 355] |  |
| **Vital status** |  |  |  |
| Living | 312 (52.4%) | 483 (37.3%) | <0.001* |
| Deceased | 283 (47.6%) | 813 (62.7%) |  |

**Notes:** \*P < 0.05.
**Abbreviations:** JAM2, The junctional adhesion molecule B; TCGA, The Cancer Genome Atlas; ER, estrogen receptor; PR, progesterone receptor; HER2, human epidermal growth factor receptor 2.

**Fig 2.**
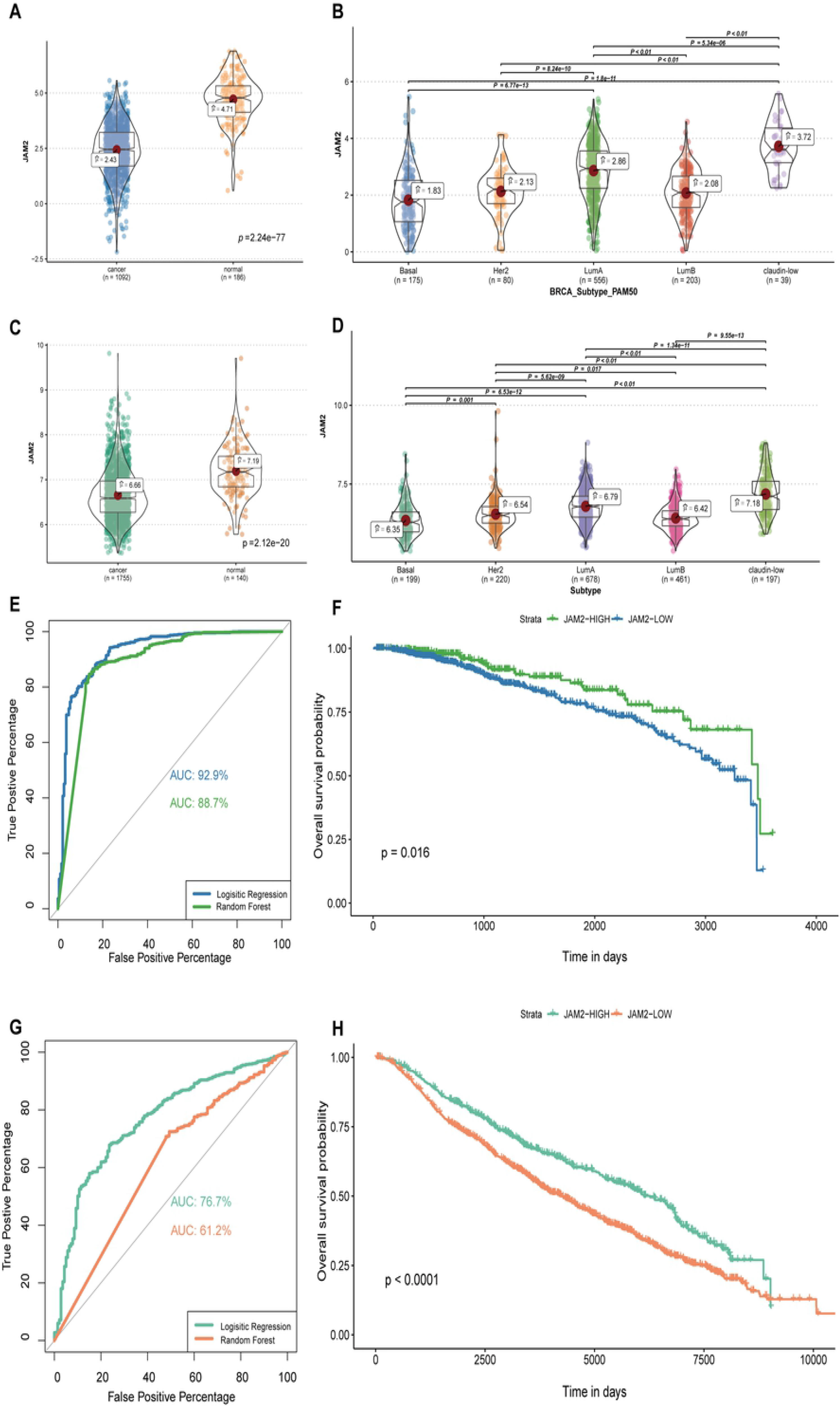
The expression and prognostic value of JAM2 in BRCA tissues and normal tissues compared by bioinformatic analysis in different databases. JAM2 mRNA is expressed at low levels in BRCA tissues (A) and differentially expressed in BRCA subtypes (B) in the TCGA cohort. JAM2 expression in BRCA is compared with that in normal samples (C) and BRCA subtypes (D) in the METABRIC database. The ROC curve of JAM2 between nontumor samples and tumor samples in the TCGA cohort (E). The overall survival (OS) analysis of JAM2 expression in the TCGA cohort (F). The ROC curve of JAM2 between nontumor samples and tumor samples in the METABRIC cohort (G). The OS analysis of JAM2 expression in the METABRIC cohort (H).

Univariate and multivariate analyses were also performed to identify the prognostic value of JAM2. Multivariate Cox regression analysis was conducted by utilizing the significant prognostic factors identified in the univariate analysis (P < 0.05). The expression of JAM2 was determined to be an independent prognostic factor (hazard ratio (HR) = 0.62, P = 0.034) in the TCGA cohort (Fig 3A). Furthermore, tumor stage (HR = 3.06, P < 0.01) and age (HR = 2.29, P < 0.01) were also independent prognostic predictors in the TCGA cohort (Fig 3A). The ER, PR and HER2 statuses were also associated with BRCA prognosis; however, there were too many patients with incomplete in the TCGA cohort. Therefore, the METABRIC dataset, which included BRCA patients and comprehensive clinical information, was used to validate the predictive prognostic value of JAM2. In the METABRIC cohort, after taking into account factors such as ER, PR and HER2 statuses, high JAM2 mRNA expression (HR = 1.88, P = 0.004) was also an independent prognostic factor by multivariate analysis (Fig 3B).

**Fig 3.**
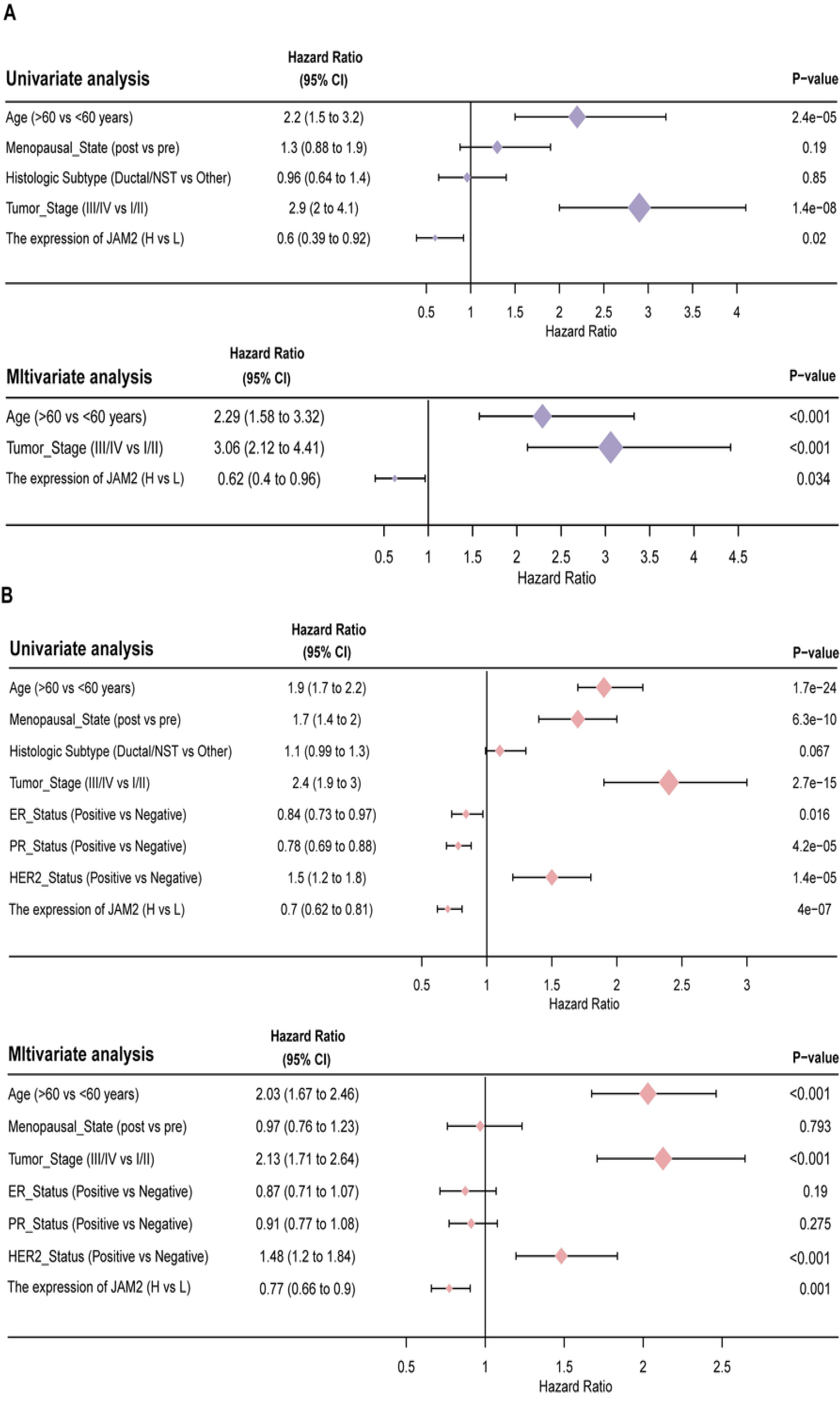
Univariate and multivariate regression analyses of the relationship between the expression of JAM2 and clinical factors in the TCGA cohort (A) and METABRIC cohort (B). Multivariate Cox regression analysis was conducted by utilizing the significant prognostic factors identified in the univariate analysis (P < 0.05).

### Hypermethylation of JAM2 in BRCA

To identify the downregulated mechanism of JAM2 in BRCA, multiple methods were used to analyze the methylation status. Three methods, the beta value of the probe with the largest standard deviation (Fig 4A), the mean beta value of all probes within the gene (Fig 4B) and the beta value of the probe with the smallest correlation coefficient with mRNA expression (strongest negative correlation) (Fig 4C), were used to represent the methylation levels of JAM2, and all methods demonstrated that JAM2 was significantly more highly methylated in BRCA tissues than in normal tissues (P < 0.05, Fig 4A, 4B and 4C). Pearson correlation analysis of 4 DNA methyltransferases showed that the expression of JAM2 was negatively related to the expression of DNMT1 (r = −0.12, P <0.01), DNMT3B (r = −0.14, P <0.01) and EHMT2 (r = −0.19, P <0.01) (Fig 4D-G). In addition, the high beta value of the cg01975706 probe, which was the most negatively correlated with the expression of JAM2 (Fig 4H), indicated a worse survival rate (Fig 4I). The above information suggests that the low expression of JAM2 in BRCA may be due to hypermethylation.

**Fig 4.**
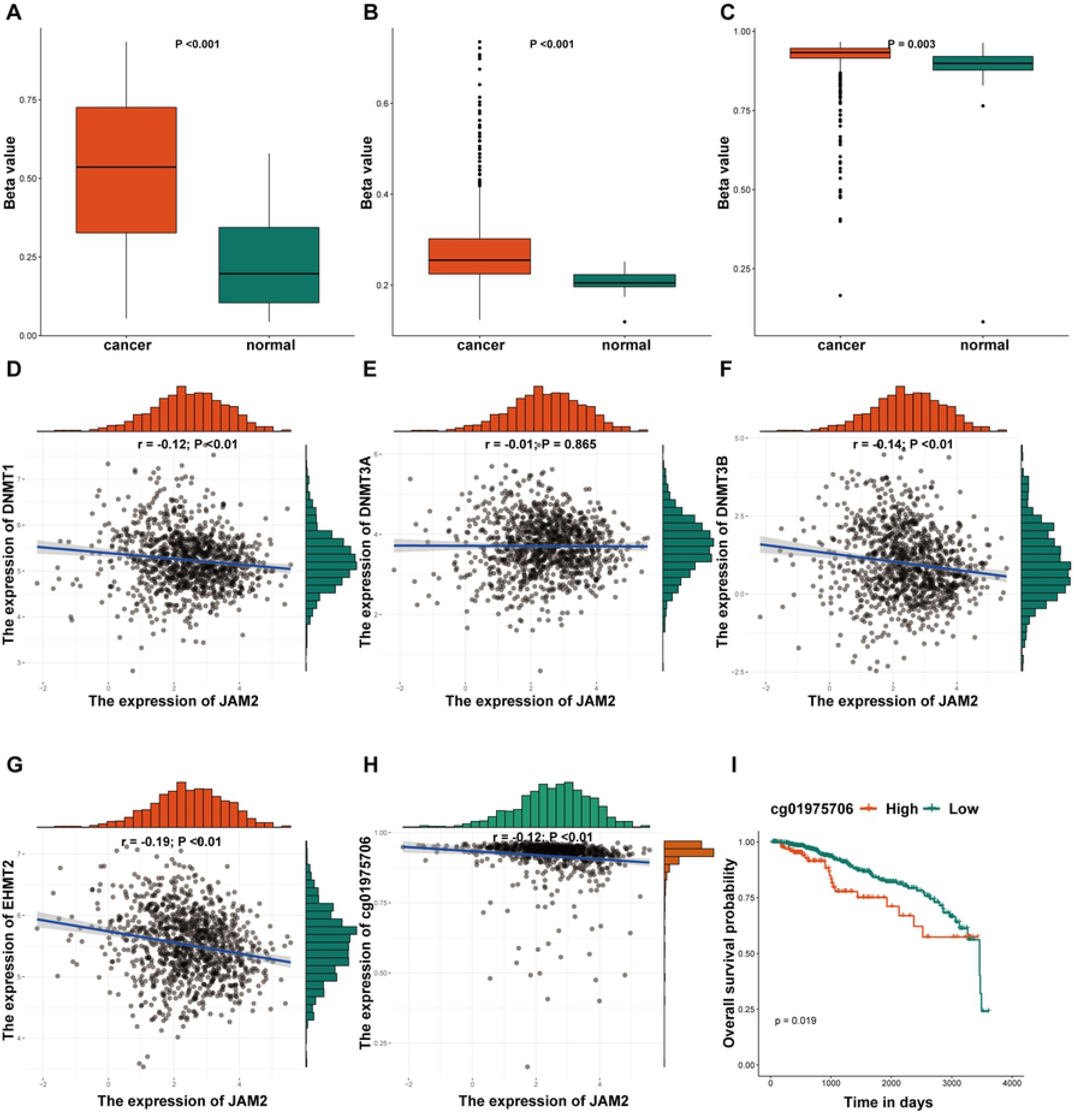
The methylation analysis of JAM2. The beta value of the probe with the largest standard deviation (A), the mean beta value of all probes within the gene (B), and the beta value of the probe with the smallest correlation coefficient with mRNA expression (strongest negative correlation) (C) represent the methylation levels of JAM2 between cancer and normal tissues. Correlation analysis between the expression of JAM2 and DNMT1 (D), DNMT3A (E), DNMT3B (F), EHMT2 (G), and the cg01975706 probe (H). Survival analysis of cg01975706 in terms of OS (I).

### Exploring potential molecular mechanisms of JAM2 in BRCA

To identify the potential molecular mechanisms of JAM2 in BRCA, 694 oncogenes were obtained from the *ONGene* database (http://ongene.bioinfo-minzhao.org/), 16 of which were associated with OS (P <0.05) and positively related to JAM2 (r > 0.4; P < 0.01) in the TCGA cohort (Fig 5A, 5B). Furthermore, 589 tumor suppressor genes (TSGs) were downloaded from the TSGene 2.0 database (https://bioinfo.uth.edu/TSGene/), and 17 TSGs associated with OS were strongly associated with JAM2 (Fig 5C, 5D).

**Fig 5.**
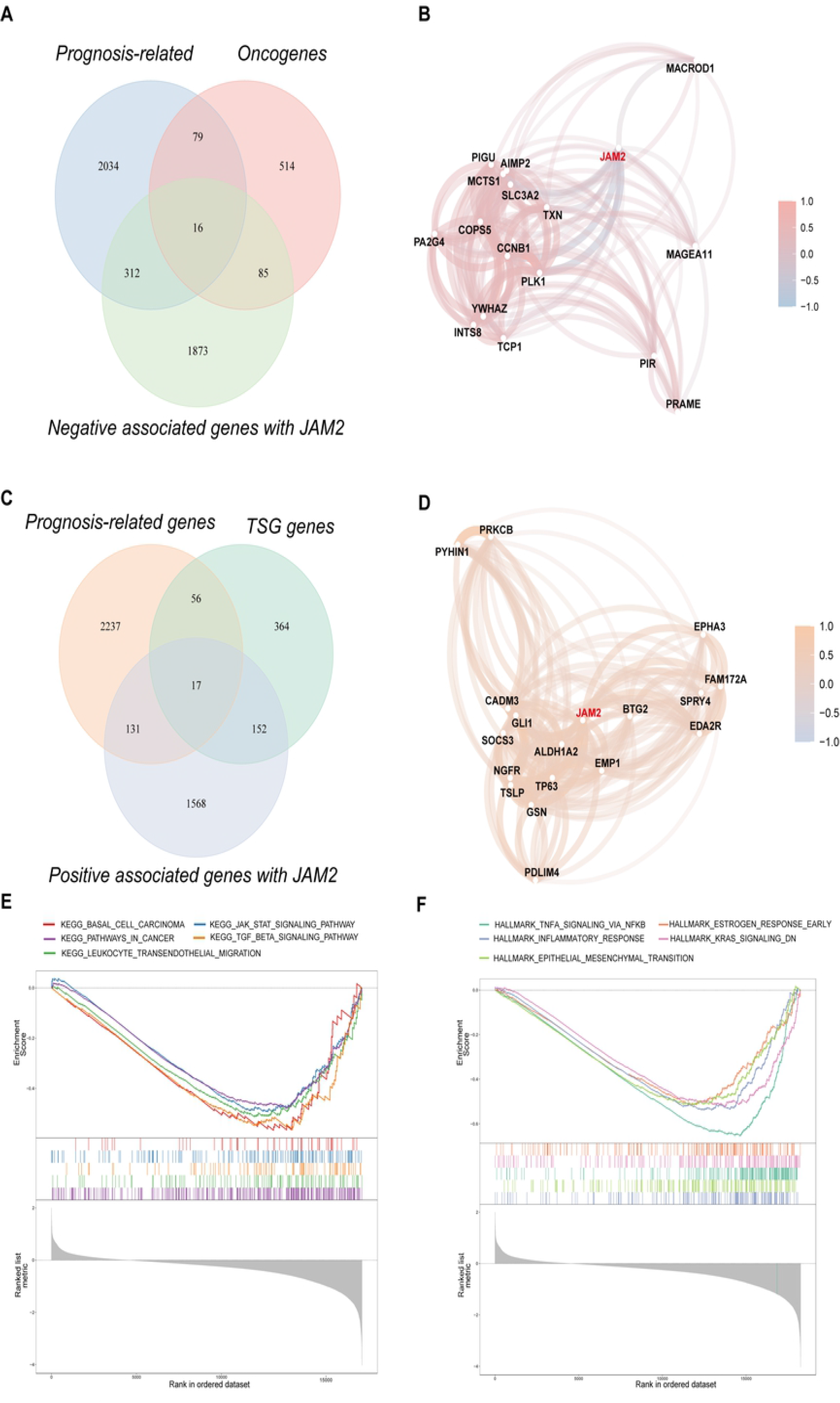
Exploring potential molecular mechanisms of JAM2 in BRCA. The intersection of 694 oncogenes was obtained from the ONGene database; 1868 genes were negatively related to JAM2 (r <0; P < 0.01), and 2441 genes were associated with OS (P <0.05) in the TCGA cohort (A). The correlation network between the 16 oncogenes and JAM2 (B). The intersection of 589 TSGs was obtained from the TSGene 2.0 database; 2286 genes were positively related to JAM2 (r > 0.4; P < 0.01), and 2441 genes were associated with OS (P <0.05) in the TCGA cohort (C). The correlation network between the 17 TSGs and JAM2 (D). GSEA revealed significant differences in the enrichment of (E) c2.cp.kegg.v7.1.entrez.gmt and (F) h.all.v7.1.entrez.gmt.

GSEA was conducted to identify the potential biological mechanisms of the JAM2 high expression group and the JAM2 low expression group in BRCA. The MSigDB results of c2.cp.kegg.v7.1.entrez.gmt indicated that the gene signatures of “pathways in cancer”, “leukocyte transendothelial migration”, “TGF beta signaling pathway”, “jak/stat signaling pathway”, and “basal cell carcinoma” were significantly suppressed in the JAM2 high expression group (Fig 5E) (Table S2). The MSigDB results of h.all.v7.1.entrez.gmt indicated that the gene signatures of “inflammatory response”, “TNFA signaling via NFKB”, “epithelial mesenchymal transition”, “KRAS signaling down”, and “estrogen response early” were significantly suppressed in the JAM2 high expression group (Fig 5F) (Table S3). Overall, these findings strongly indicate the potential role of JAM2 as a tumor suppressor in BRCA.

### The use of reverse-phase protein array (RPPA) to explore the potential function of JAM2 in BRCA

The RPPA pathway scores, which correlate with tumor lineage and indicate the protein expression signatures of pathway activity, were obtained by previously published literature in TCGA[24]. The analysis of the correlation between the expression of JAM2 and the pathway scores of RPPA indicated that JAM2 was most positively related to the epithelial-mesenchymal transition (EMT) score (r = 0.38; P <0.01) and most negatively associated with the proliferation score (r = −0.56; P <0.01) (Fig 6A). In addition, all pathway scores were significantly related to the expression of JAM2 except for the hormone a score, apoptosis score and PI3K/Akt score (Fig 6A) (Table S4). Our analysis also indicated that the pathway scores for EMT, Ras/MAPK and RTK were significantly higher in the high JAM2 expression group. However, the pathway scores for tumor purity, proliferation, cell cycle, DNA damage response and TSC/mTOR were significantly lower in the high JAM2 expression group (Fig 6B–I). Furthermore, correlation analysis was also performed between the expression of JAM2 and 226 tagged proteins in RPPA. The results indicated that 59 proteins were positively correlated with JAM2 (P < 0.01) and 72 proteins were negatively correlated with JAM2 (P < 0.01) (Fig 7A) (Table S5). The three proteins most positively and negatively associated with JAM2 are shown in Fig 7 B-G.

**Fig 6.**
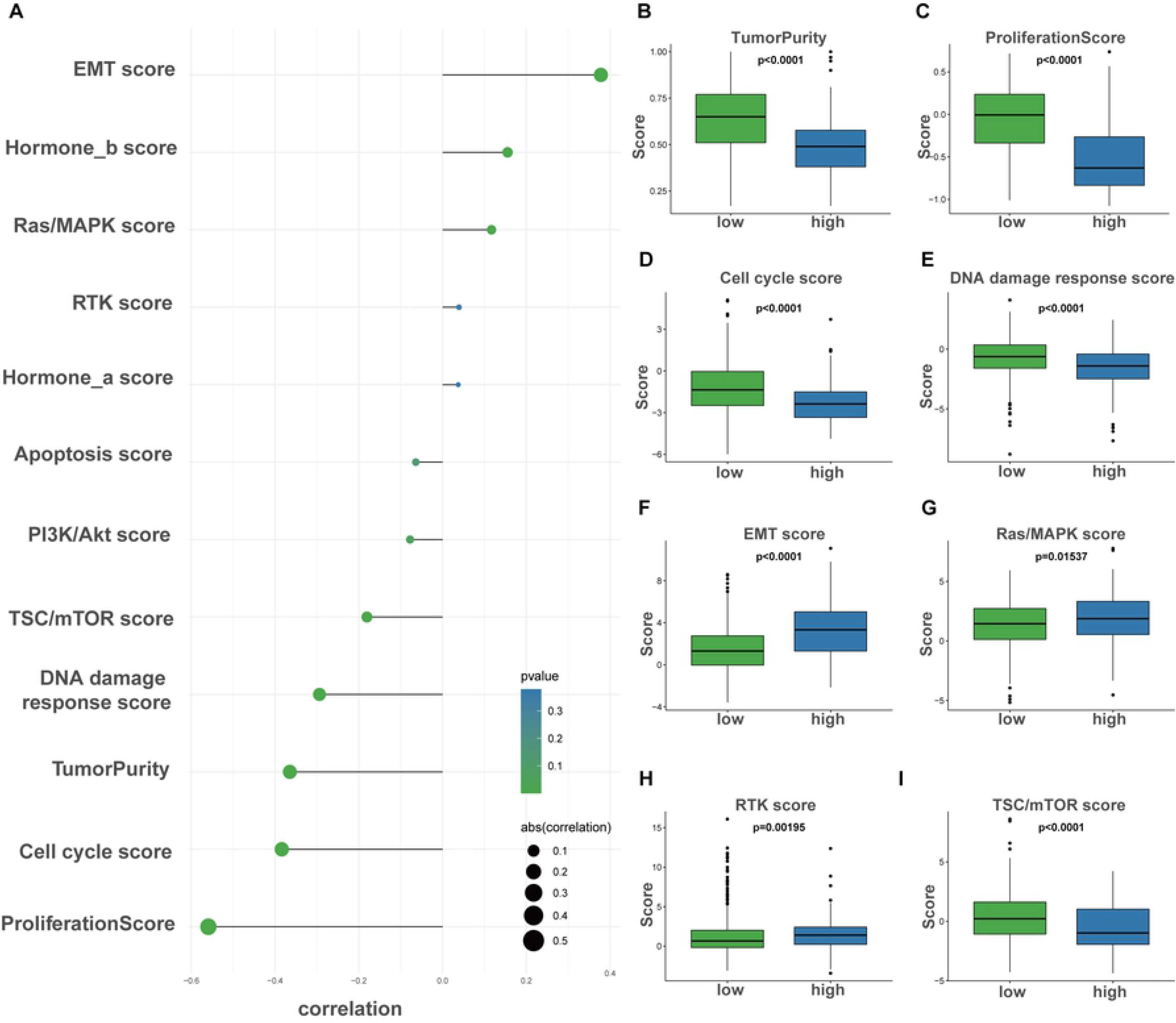
Phenotype heterogeneity between different JAM2 expression groups. The correlation analysis bubble diagram shows the relationship between the different RPPA pathway scores and the expression of JAM2 (A). Different colors represent different p-values, and the size of the circle represents the correlation coefficient. Boxplots show differences in (B) tumor purity, (C) proliferation, (D) cell cycle, (E) DNA damage response, (F) EMT, (G) Ras/MAPK, (H) RTK and (I) TSC-mTOR scores between groups with high and low JAM2 expression in the TCGA cohort.

**Fig 7.**
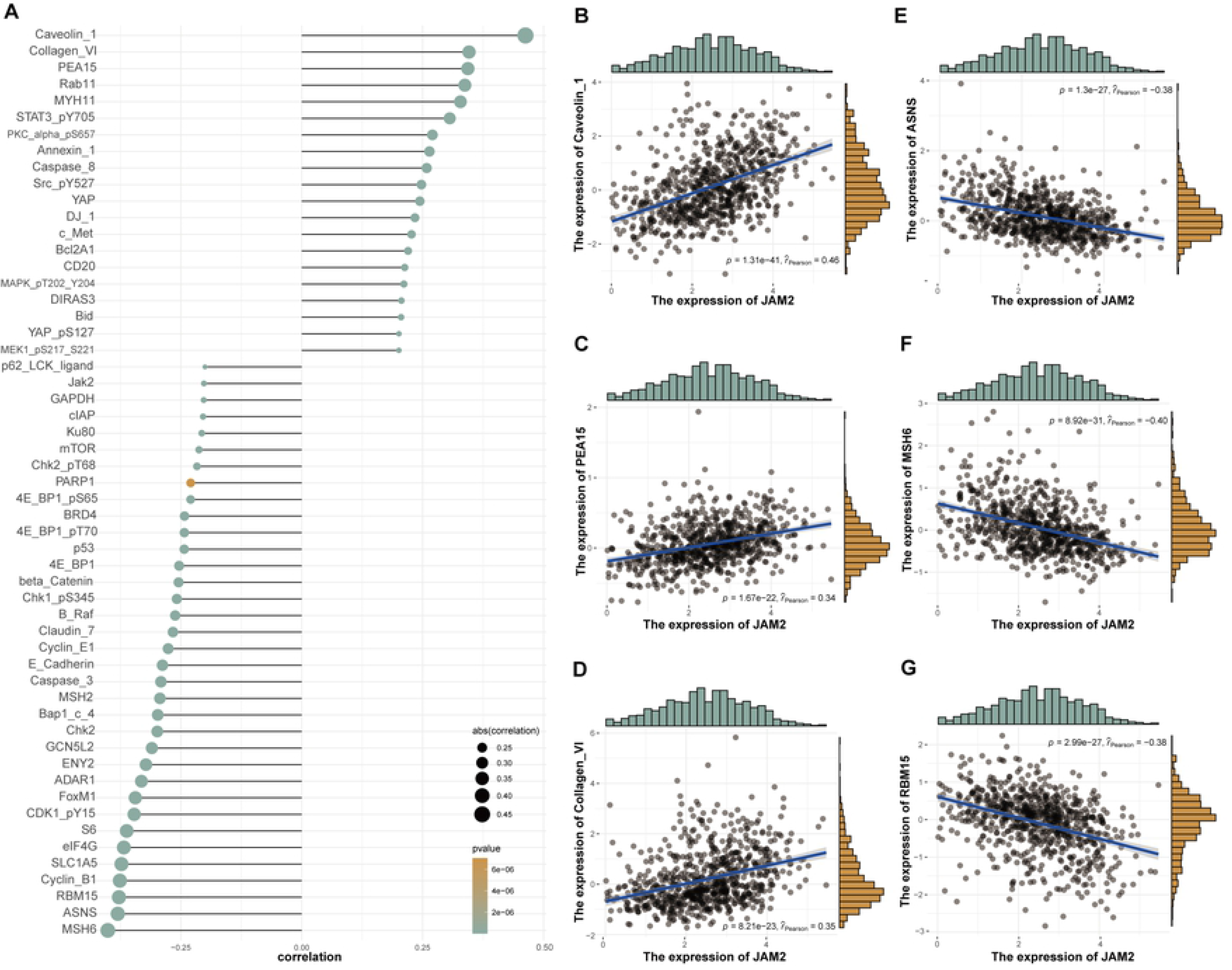
Correlation analysis between the expression levels of JAM2 and 226 tagged proteins in RPPA. The correlation analysis bubble diagram shows that 59 proteins were positively correlated with JAM2 (P < 0.01) and 72 proteins were negatively correlated with JAM2 (P < 0.01) (A). Different colors represent different p-values, and the size of the circle represents the correlation coefficient. The three proteins most positively associated with JAM2 expression are Caveolin 1 (r = 0.46; P <0.01), Collagen VI (r = 0.35; P <0.01) and PEA15 (r = 0.34; P <0.01) (B-G); the three proteins most negatively related with JAM2 expression are CMSH6 (r = −0.40; P <0.01), ASNS (r = −0.38; P <0.01) and PEA15 (r = −0.38; P <0.01) (B-G).

### Relationships between JAM2 expression and neoantigens, tumor mutational burden (TMB) and microsatellite instability (MSI)

Most cancers are associated with mutations and genomic instability, such as chromosomal instability (CIN) or MSI[25]. Neoantigens are nascent antigens encoded by mutant genes in tumor cells, mainly produced by point mutations, deletion mutations, gene fusions, etc., that are not the same as proteins expressed by normal cells[26]. Neoantigens can be utilized to identify tumor-specific T cells or to generate vaccines[27]. The TMB was estimated by analyzing somatic mutations, including single-nucleotide variants (SNVs) and insertions/deletions (INDELs) per megabase of the panel sequences examined, and can be a potential predictive biomarker for immunotherapy[28]. Therefore, the MSI scores, number of neoantigens and TMB in each TCGA sample were calculated, and significant negative correlations were found between JAM2 expression and MSI (r = −0.08; P = 0.013), neoantigens (r = −0.18; P < 0.001) and TMB (r = −0.09; P = 0.007) (Fig 8A-C).

**Figure 8.**
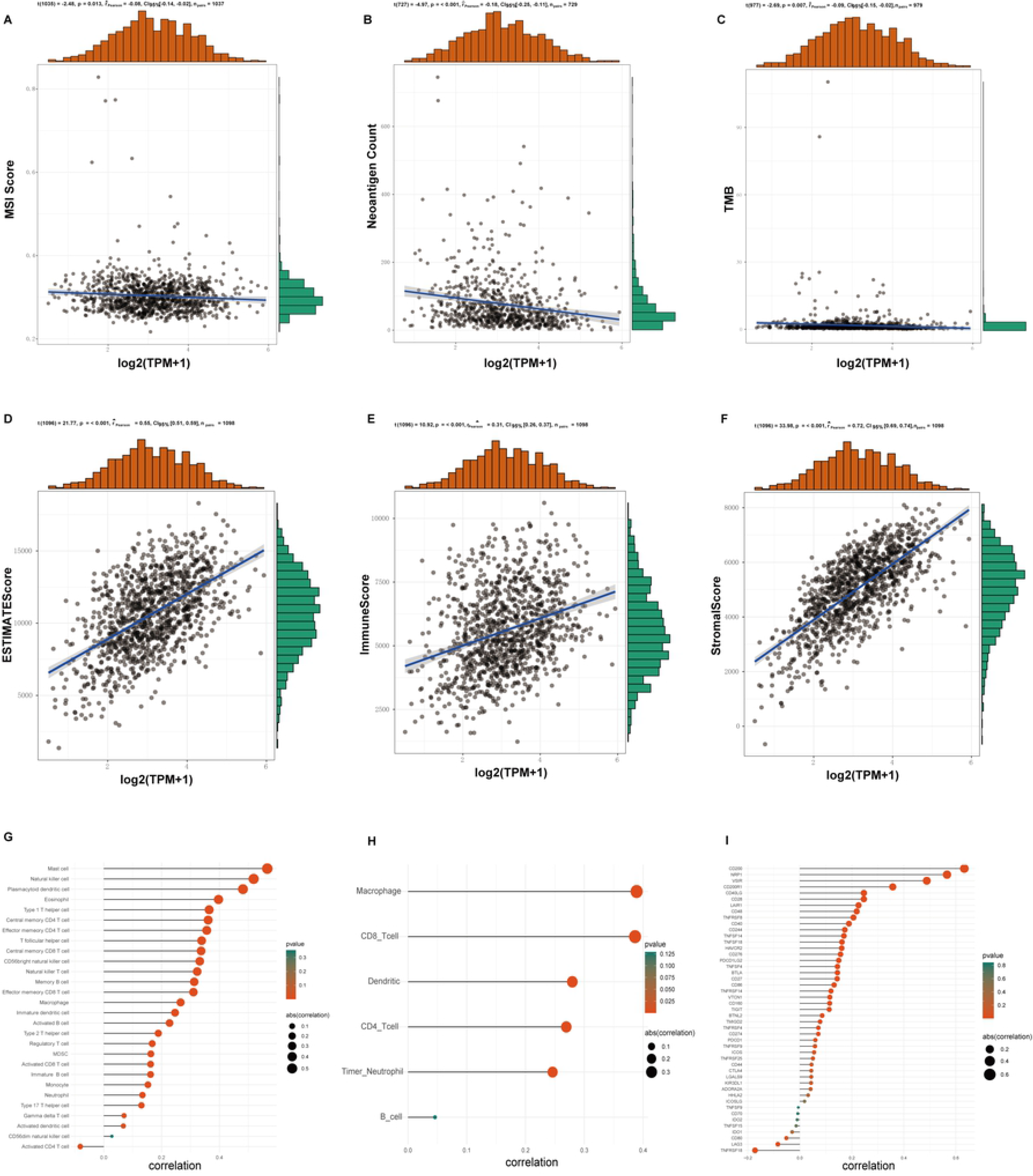
Correlation analysis of immune-related scores and JAM2 expression. The correlation analysis indicated that the MSI score (r = −0.08; P = 0.013) (A), neoantigen counts (r = −0.18; P <0.001) (B) and TMB (r = −0.09; P = 0.007) (C) were negatively correlated with the expression of JAM2. The correlation analysis showed that the estimate score (r = 0.55; P = 0.013) (D), immune score (r = 0.31; P < 0.001) (E) and stromal score (r = 0.72; P < 0.001) (F) were positively correlated with the expression of JAM2. The correlation analysis bubble diagram shows the relationships among the 28 immune infiltration cell scores from TCIA (G), 6 immune infiltration cell scores from TIMER (H) and the expression of JAM2. The correlation bubble diagram shows the relationship between common immune checkpoint genes and JAM2 (I). TCIA = The Cancer Immunome Atlas.

### Immunotherapy and chemotherapy responses of high- and low-JAM2 patients in BRCA

Increasing reports indicate that the tumor immune microenvironment plays an important role in tumor development. Therefore, the estimate scores, immune scores and stromal scores were analyzed by the *estimate* R package. The correlation analysis indicated that the expression of JAM2 was positively associated with these genes (all P <0.05) (Fig 8D-F). Tumor-infiltrating lymphocytes are independent predictors of lymph node status and survival in cancer patients. Twenty-eight immune infiltration cell scores and 6 immune infiltration cell scores were downloaded from The Cancer Immunome Atlas (https://tcia.at/)[29] and Tumor IMmune Estimation Resource (TIMER)[30], respectively. The results indicated that except for the CD4 T cell score, all immune infiltration cell scores were positively correlated with JAM2 expression (Fig 8G-H). Common immune checkpoint genes were also collected to analyze the relationship between JAM2 expression and immune checkpoint gene expression. The correlation bubble diagram showed that most of the immune checkpoint genes were significantly positively correlated with JAM2 (Fig 8I).

Immunotherapy using immune checkpoint blockade targeting CTLA-4 and PD-1 has emerged as a promising strategy for the treatment of various malignancies[31]. Therefore, ImmuCellAI[22], which is a gene set signature‐based method, was used to estimate the clinical response to immune checkpoint blockade in the TCGA and METABRIC cohorts. The results indicated that JAM2 expression was higher in those predicted to be sensitive to immunotherapy than in those who were not sensitive (Fig 9A, 9C), and the immune infiltration scores were also different between the high and low JAM2 expression groups in the TCGA and METABRIC cohorts (Fig 9B, 9D).

**Figure 9.**
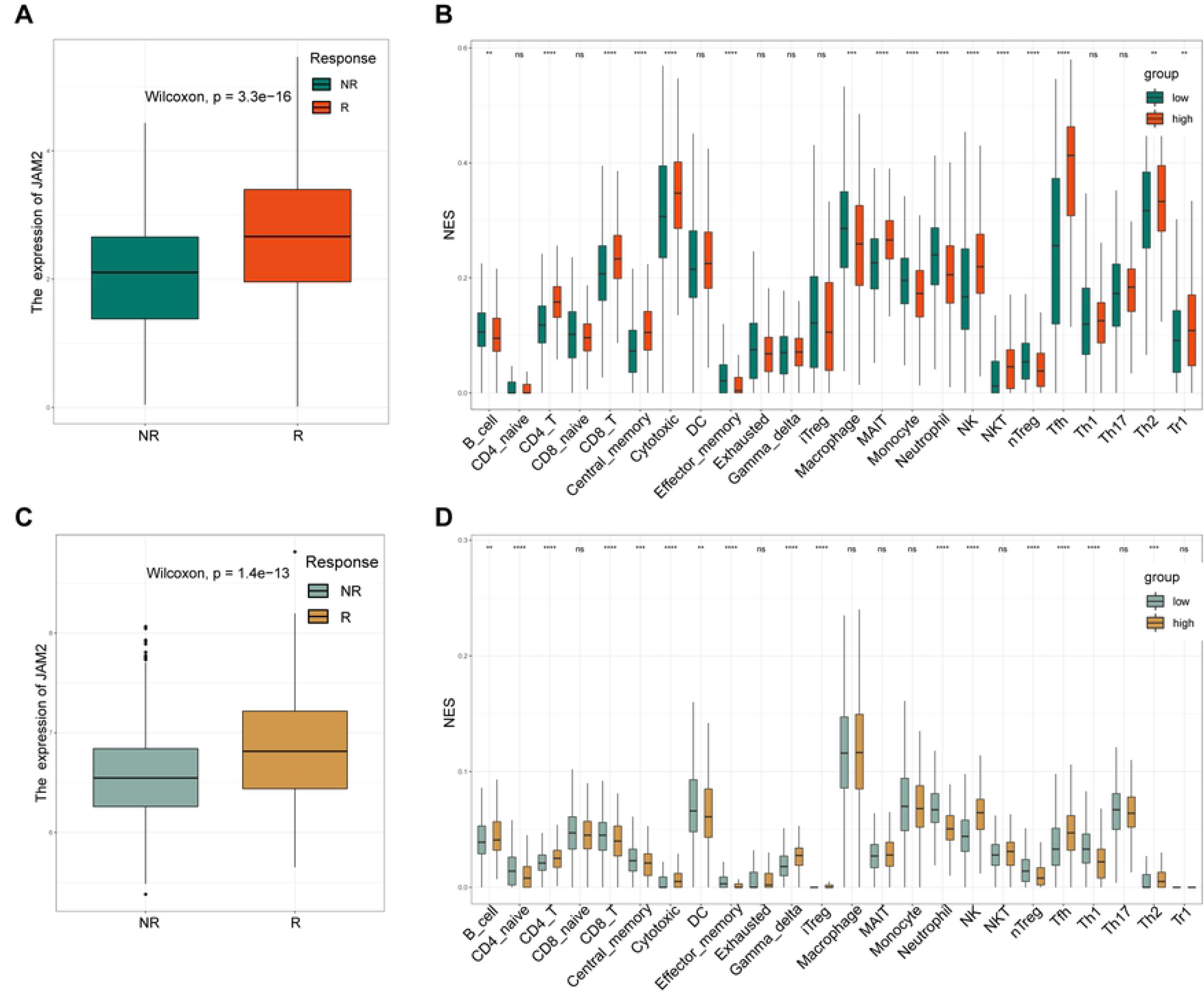
The immunotherapeutic responses to anti-CTLA-4 and anti-PD-1 treatments of patients with different JAM2 expression levels in the TCGA cohort (A) and the METABRIC cohort (C). The immune infiltration scores of high- and low-JAM2 expression patients in the TCGA cohort (B) and the METABRIC cohort (D).

The IC50 value of each BRCA patient in the TCGA cohort was predicted by the *pRRophetic* R package[23]. BRCA patients with a low expression of JAM2 have a worse prognosis. The 18 chemotherapy drugs that patients in the JAM2 low expression group were more sensitive to are shown in Fig 10 (Table S6). These results may provide clinical treatment strategies for future studies.

**Figure 10.**
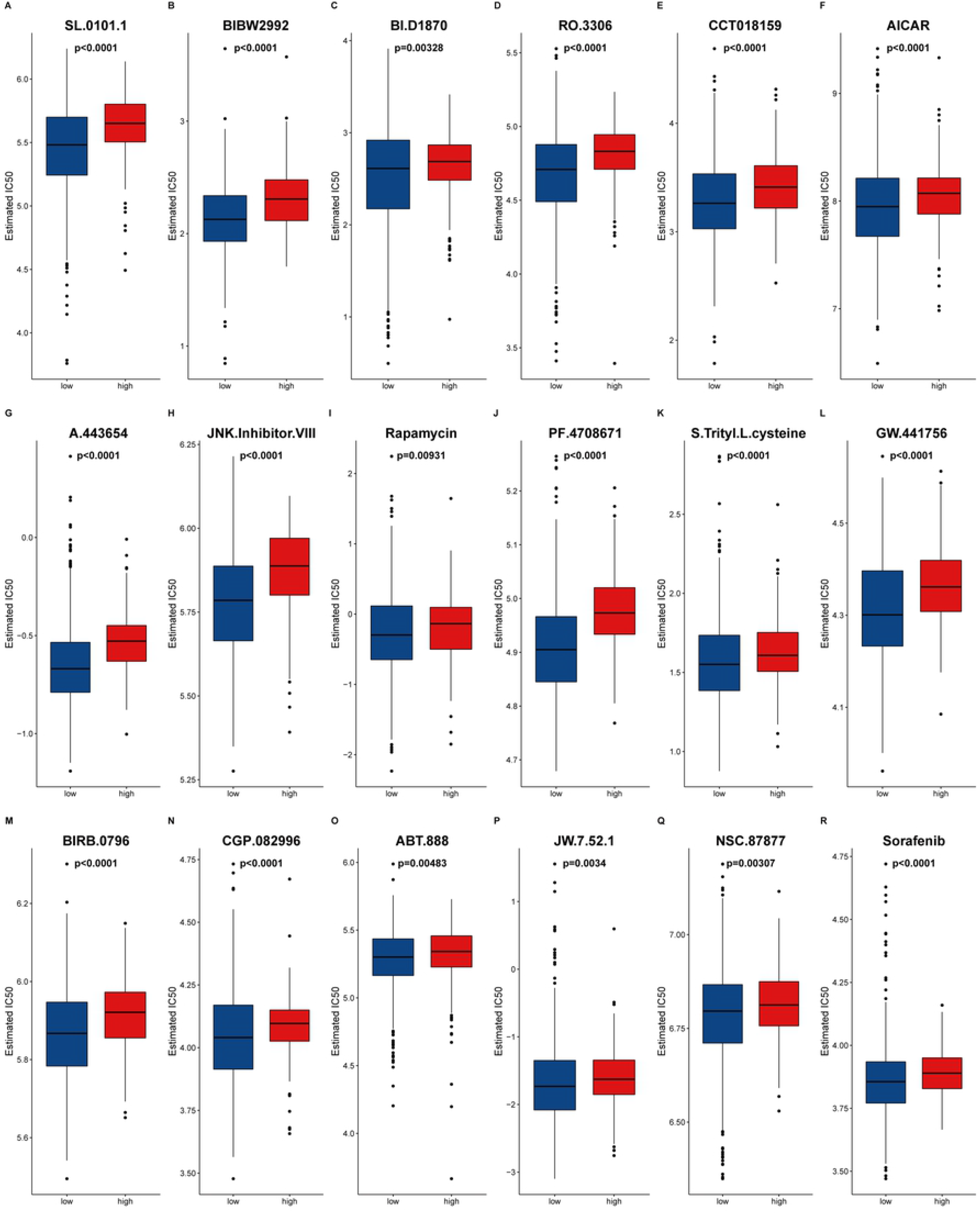
The 18 chemotherapy drugs that showed more sensitivity in the JAM2 low expression group (A - R).

## Discussion

JAM2 belongs to a subfamily of immunoglobulins that regulate tight junction assembly and leukocyte recruitment during inflammation, angiogenesis, cell migration and proliferation[32]. More research has shown that JAM2 is differentially expressed in different types of cancers. Kok-Sin et al found that the expression of JAM2 was downregulated due to its hypermethylated status in colorectal cancer[33]. Similarly, the methylation of JAM2 was increased, and its expression was decreased in esophageal squamous cell carcinoma[34]. However, in some tumors, JAM2 seems to play a different role. Arcangeli et al showed that JAM-B expressed by endothelial cells is involved in the metastasis of melanoma cells through the interaction with JAM-C on the tumor[35]. Lifeng Qi et al indicated that the deregulation of JAM2 in glioma cell lines can lead to profoundly decreased proliferation and migration of cells in vitro[36]. Hajjari et al found that JAM2 was significantly upregulated in gastric tumor samples compared with adjacent normal tissues[15].

In our study, multiple databases were used to explore the potential role and underlying mechanism of JAM2 in BRCA. The expression of JAM2 was relatively low in BRCA cell lines compared to other cancer cell lines from the CCLE database (Fig 1A), and the expression of JAM2 was relatively high in normal breast tissue compared to other normal tissues in the GTEx database (Fig 1B). Interestingly, except for cholangiocarcinoma (CHOL), glioblastoma multiforme (GBM), acute myeloid leukemia (LAML), low-grade glioma (LGG), liver hepatocellular carcinoma (LIHC) and pancreatic adenocarcinoma (PAAD), downregulation was observed in all other tumors compared with that in normal tissues through analyses of the GTEx and TCGA databases (Fig 1C).

DNA methylation is an imperative epigenetic mechanism that regulates several biological processes. Aberrant DNA methylation in cancer is typically characterized by the promoter-specific hypomethylation of genes, particularly cancer suppressor genes, leading to gene silencing[37, 38]. Therefore, the methylation status of JAM2 in BRCA was analyzed by multiple methods to identify the downregulated mechanism of JAM2 in BRCA. Finally, similar to that in esophageal squamous cell carcinoma and colorectal cancer, we found that the expression of JAM2 was downregulated, which may be due to the hypermethylated status of the JAM2 gene in BRCA (**Fig 4A-4I**).

The role of JAM2 in tumorigenesis and development has also been investigated in previous research. Huang JY et al suggested that JAM2 and JAM-C may potentially be involved in gastric adenocarcinoma tumor progression[39]. Qi LF also observed that downregulated JAM2 could inhibit glioma migration in vitro by blocking the NOTCH pathway[40]. An interesting finding suggested that overexpression of JAM-B is responsible for suppressing VEGF-induced angiogenesis in a Down syndrome trisomy 21 mouse model, thereby restraining the tumor effects in a mouse model of lung cancer[17]. JAM2 is involved in BRCA progression, but the underlying mechanism remains unclear[41]. To identify the potential molecular mechanisms of JAM2 in BRCA, 16 oncogenes and 17 TSGs associated with OS and JAM2 were identified. GSEA showed that the gene signatures of “epithelial mesenchymal transition”, “pathways in cancer”, “jak/stat signaling pathway” and “basal cell carcinoma” were significantly suppressed in the JAM2 high expression group. These findings strongly indicate the potential role of JAM2 as a tumor suppressor in BRCA.

The analysis of proteomic data is one of the commonly used tools in the field of tumor pathophysiology, and the RPPA technique is widely used in the TCGA project to study the tumor proteome[24]. The three proteins most positively and negatively associated with JAM2 were caveolin 1, collagen VI, and PEA15 and MSH6, ASNS, and RBM15. Furthermore, Ludwig et al found that JAM2 plays a role in T lymphocyte rolling and firm adhesion by interacting with α4β1[42]. Therefore, the estimate scores, immune scores and stromal scores were analyzed by the estimate R package, and the expression of JAM2 was positively associated with the abovementioned scores (all P <0.05). In addition, except for the CD4 T cell score, all immune infiltration cell scores were positively correlated with JAM2 expression in different databases. We estimated the clinical response to immune checkpoint blockade in the TCGA and METABRIC cohorts and found that JAM2 expression was higher in those predicted to be sensitive to immunotherapy than in those who were not sensitive. These results suggest that JAM2 plays a broad role in BRCA inflammatory infiltration and that high expression of JAM2 may predict a better immunotherapy response.

Finally, the *pRRophetic* R package was used to predict the IC50 of each BRCA patient in the TCGA cohort. Eighteen chemotherapy drugs that patients in the JAM2 low expression group were more sensitive to were found to provide clinical treatment strategies in the following study.

## Conclusions

Our results demonstrated the diagnostic and prognostic value of JAM2. Analysis of the molecular mechanisms indicates the potential role of JAM2 as a tumor suppressor, and high JAM2 expression may predict a better immunotherapy response in BRCA.

## Data Availability

The datasets used and/or analyzed during the current study are available from the corresponding author on reasonable request. We downloaded 21 tumor cell lines from the Cancer Cell Line Encyclopedia (CCLE) database (https://portals.broadinstitute.org/ccle/data) and 31 normal tissues from the Genotype-Tissue Expression (GTEx) database (http://www.gtexportal.org). The pan-cancer‐normalized data of 27 tumor and normal tissues were downloaded from UCSC Xena (https://tcga.xenahubs.net). The clinical and RNA sequencing (RNA-seq) data of 947 patients in The Cancer Genome Atlas (TCGA) cohort and 1891 patients in the Molecular Taxonomy of Breast Cancer International Consortium (METABRIC) cohort of BRCA were obtained from TCGA portal (https://tcga-data.nci.nih.gov/tcga/) and cBioPortal (http://www.cbioportal.org). Data on the methylation status of BRCA were extracted from the FireBrowse database (http://www.firebrowse.org).

## Acknowledgements

Not applicable.

## Author Contributions

Conceived and designed the experiments: CQ and YP. Performed the experiments: YZZ and BGZ. Analyzed the data: YF, JX. Contributed reagents/materials/analysis tools: SCL. Wrote the paper: CQ and JX.

